# Cerebellar involvement in self-timing

**DOI:** 10.1101/2025.03.05.641568

**Authors:** Ellen Boven, Jasmine Pickford, Richard Apps, Nadia L Cerminara

## Abstract

The cerebellum is well-established in sub-second motor timing, but its role in supra-second interval timing remains unclear. Here, we investigate how cerebellar output influences time estimation over longer timescales. Rats performed an interval timing task, estimating time based on an auditory cue, while chemogenetic inhibition of the lateral cerebellar nucleus assessed its role in both predictable (externally cued) and unpredictable (internally cued) timing conditions. Cerebellar inhibition produced bidirectional effects: delayed action initiation in predictable trials and premature responses in unpredictable trials. Despite slowed movement, overall task success rates remained unchanged, suggesting a specific impairment in temporal estimation rather than motor execution. These findings demonstrate that the cerebellum integrates motor and cognitive processes for supra-second timing, with differential effects on externally guided and self-generated timing. Our results provide evidence that the lateral cerebellum contributes to supra-second interval timing, supporting its role in adaptive behavior across extended timescales.

## Introduction

The ability to produce sequential movements underlies everyday behaviour, with timing serving as avital component in motor control (Avanzino et al., 2016). External stimuli provide temporal cues to enable coordination with moving objects in dynamic environments, while an internal perception of time intervals is required when participants are required to estimate time duration and execute a task or compare the duration to a reference duration (Grondin, 2010). Self-timing – the ability to initiate movements based on an internally generated temporal framework rather than external sensory cues, is a key aspect of this process.

Timing task scan also be divided by their temporal scale (Buhusi and Meck, 2005). Sub-second timing, which operates within milli second intervals, is crucial for rapid motor functions like locomotion, reflexes, eye movement and object manipulation – functions that are dependent on an intact cerebellum (Garcia& Mauk,1998; Krupa& Thompson, 1997; Mojtahedian et al., 2007). Conversely, supra-second timing which operates over intervals ranging from seconds to minutes, is associated with goal-directed behaviours requiring higher cognitive processes such as foraging and decision making (Pezzulo et al., 2014; Gupta, 2014; Friendman & Robbins, 2022; Szelag et al, 2022). While much is known about cerebellar contributions to sub-second timing (for reviews see Ivry et al., 2002; Breska & Ivry, 2016; Tanaka et al., 2021; Boven & Cerminara, 2023) its contributions to supra-second timing remain poorly understood.

Human studies, including cerebellar patients and functional brain imaging, have demonstrated a possible role for the cerebellum insupra-secondtiming (Nichelli et al., 1996; Malapani et al., 1998; Mangels et al., 1998; Tracy et al., 2000;O’Reilly et al., 2008; Gooch et al., 2010). Supra-second timing is often studied through behavioural paradigms of interval timing, which require individuals to monitor time over several seconds to minutes to guide goal-directed actions and decision-making. These behaviours involve cognitive components, where individuals estimate and act on time intervals that are longer than those typically studied in cerebellar research, yet have been instrumental in demonstrating a potential role for the cerebellum in supra-second timing. For example, Gooch et al., (2010) observed that cerebellar patients with damage to the lateral regions of the cerebellum exhibited both overestimation and underestimation of supra-second intervals during temporal estimation and production tasks. However, given the inherent methodological constraints of human studies it remains unclear how the cerebellum contributes to these behaviours.

Interval timing tasks in animals can provide insight into how the cerebellum facilitates temporal estimation while controlling for motor performance, thereby advancing our understanding of cerebellar functions across diverse timescales (Parker, 2016; Ohmae et al., 2017, Kunimatsu et al., 2018; Heskje et al., 2020; Heslin et al., 2022). Ohmae et al., (2017) and Kunimatsu et al., (2018) examined the role of the cerebellar dentate nucleus in self-timing by training monkeys to initiate saccades after delay intervals ranging from 0.4-2.4 seconds. Their findings demonstrated that preparatory neural activity in the dentate nucleus preceding self-timed saccades was temporally consistent across delay intervals, irrespective of duration, suggesting that the dentate nucleus in the cerebellum may contribute not only to sub-second timing but also to the execution and fine-tuning of movements following supra-second delays. This contrasts with findings from Heslin et al., (2022), who did not find evidence for cerebellar involvement in supra-second interval timing tasks in rodents. The discrepancy between these studies highlights ongoing uncertainty regarding the cerebellum’s role in self-timing beyond the sub-second range. Therefore, the aim of the present study was to investigate cerebellar contributions to supra-second self-timing by utilising an interval timing task in rats, while controlling for motor strategies. In this task, rats were required to estimate time based on sound duration, terminating a nose poke at the appropriate moment to receive a reward. Previous research has shown that this task recruits the prefrontal cortex but not the motor cortex (Xu et al., 2014), suggesting recruitment of higher cognitive processes in the task. Using a supra-second time estimation task and inhibitory chemogenetics in the lateral cerebellar nucleus, we show that rats underestimate time, suggesting cerebellar contributions to self-timing at supra-second intervals. These findings therefore support a role for the cerebellum in integrating motor and cognitive functions to enable adaptive behaviour over extended timescales.

## Results

### Targeting of lateral cerebellar output

Human imaging and patient studies have indicated that regions of the cerebellar hemisphere are implicated in interval timing (King et al., 2019, Gooch et al., 2010). Given that the lateral cerebellar nuclei (LCN) are the main output of the cerebellar hemispheres we targeted infusion of inhibitory Designer Receptors Exclusively Activated by Designer Drugs (DREADD) virus, AAV5-hSyn-hM4D(Gi) (termed hM4D(Gi)) or control virus AAV5-hSyn-EGFP (which lacks the DREADD receptor), into LCN bilaterally, to chronically manipulate cerebellar output during an interval timing task (Fig. 1a). Successful targeting was supported by the presence of hM4D(Gi)-mCherry or control-EGFP expressed in the LCN (Fig. 1b). While there was some expression in the cerebellar cortex this was variable across animals and was not related to any inter-animal differences in behaviour. Semi-quantitative mapping of the expression across the cerebellar nuclei (Fig. 1c, d) showed that expression was generally centred on but not restricted to the LCN.

**Figure 1.**
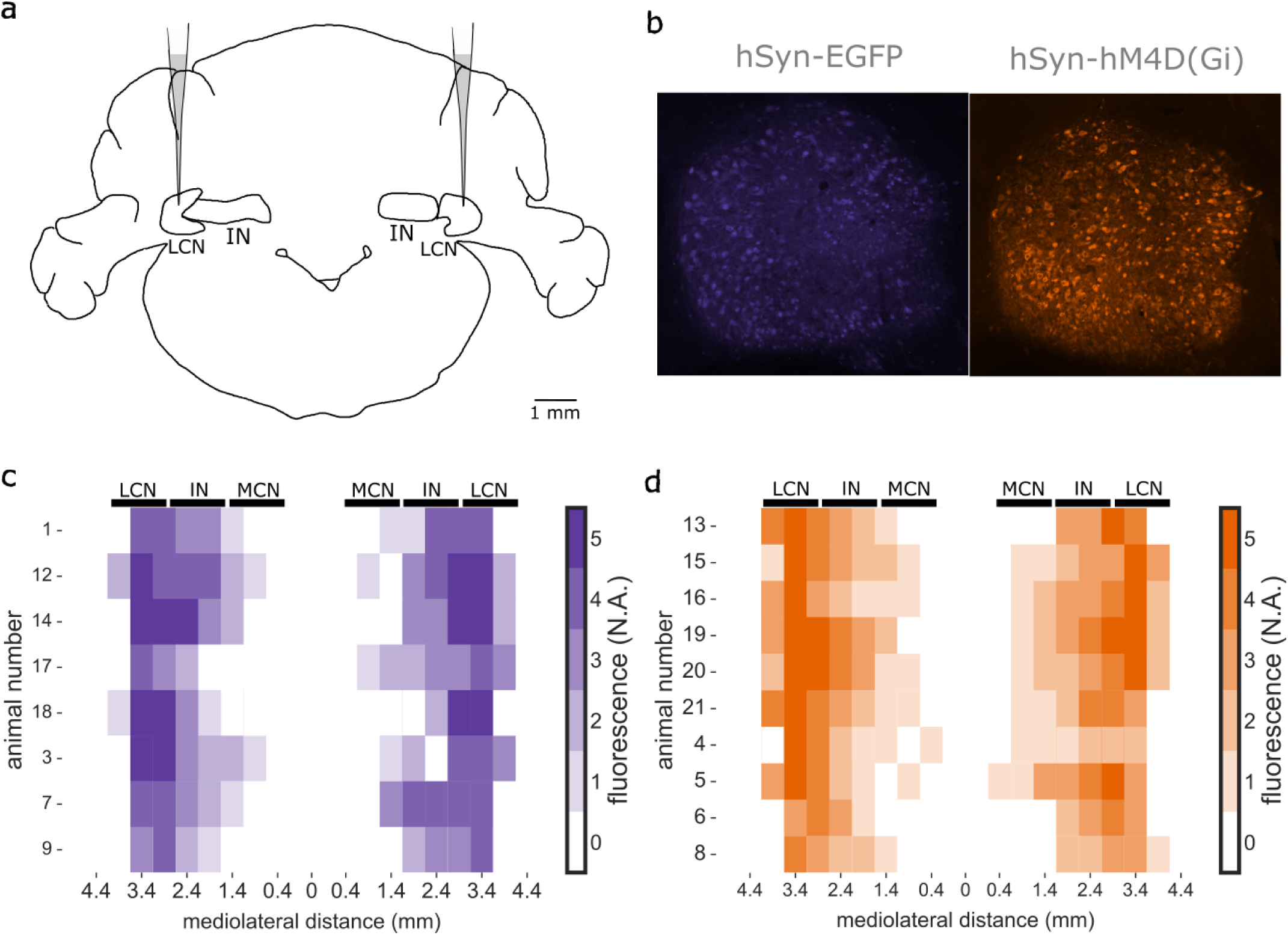
Viral expression is concentrated in LCN. (a) Schematic of viral-mediated delivery of AAV5-hSyn-EGFP or AAV5-hSyn-hM4D(Gi) bilaterally targeting the lateral cerebellar nucleus (LCN). (b) Sagittal sections of LCN showing representative examples of expression of AAV5-hSyn-EGFP and AAV5-hSyn-hM4D(Gi). (c) Heatmaps showing the extent of EGFP (n=8) expression across the mediolateral axis of the cerebellar nuclei (LCN=lateral cerebellar nucleus, IN=interpositus nucleus, MCN=medial cerebellar nucleus). Colours of heatmap indicate level of expression with no expression at 0 and maximal expression at 5. **(d)** Same as **(c)** but for hM4D(Gi) (n=10).

### Chemogenetic manipulation of cerebellar output affects general motor performance

As a first step, it was important to assess whether chemogenetic inhibition of LCN had an effect on general motor performance. To this end, we compared the effect of systemic clozapine-N-oxide (CNO) administration on open field exploration on the hM4D(Gi) group (n=6 rats) compared to the control EGFP group (n=5 rats, Fig. 2). Open field performance was assessed 30 minutes after the administration of CNO. The hM4D(Gi) group, compared to control, showed a significant reduction in open field exploration, expressed as the total distance travelled (Fig. 2a-b, hM4D(Gi) 29.45 ± 19.5 m; n=6 rats; EGFP 61.37 ± 11.71m, mean ± SD, n=5 rats: p = 0.0063, unpaired t-test); as well as a reduction in the average velocity during movement (Fig. 2c, hM4D(Gi) = 5.63 ± 3.75 cm/s, n=6 ; EGFP = 12.78 ± 2.0 cm/s, n=5, p = 0.0021). These results provide a positive control that CNO activation of DREADD transfected neurons in the cerebellar nuclei was effective but that general motor effects may be a confound in the interval timing task.

**Figure 2.**
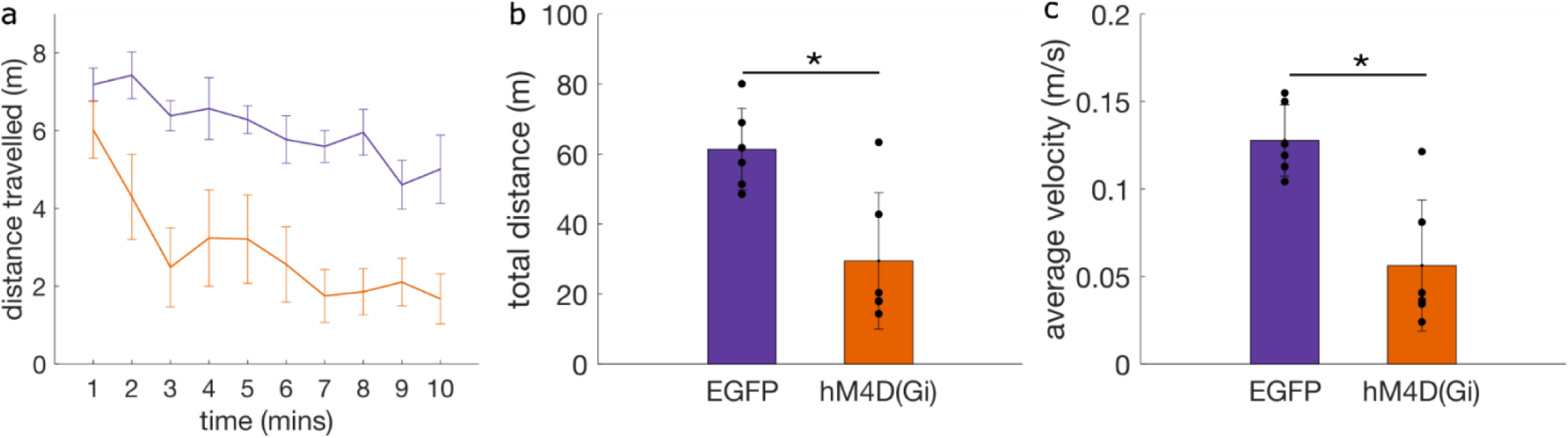
Effect of chemogenetic inhibition of cerebellar output on motor performance. (a) Average distance travelled during 10 minutes of open field exploration for EGFP (purple, n=5) and hM4D(Gi) (orange, n=6) rats per minute. (**b**) Average total distance travelled during 10 minutes of open field exploration for EGFP (n=5) and hM4D(Gi) (n=6) groups. Individual data points show total distance travelled per animal. Bars represent mean ± sem. Unpaired t-test *: *p* < 0.01 (c) Same as **(b)** but for average velocity.

### Interval timing task

To investigate how rats estimate and utilize both externally and internally cued temporal intervals to guide behaviour we used an interval timing task with predictable and unpredictable time cues(Fig. 3). Specifically, the study compared when the timing of a reward could be anticipated based on an external cue (predictable time cue, Fig. 3a) versus when the cue duration was variable and the reward timing could not be directly inferred from cue offset (unpredictable time cue, Fig. 3c). The predictable condition assessed the animals’ ability to associate a fixed 2.5-second auditory cue with a reward window, testing temporal precision in response to a consistent auditory signal (Fig. 3a). Conversely, the unpredictable condition evaluated the rats’ capacity for self-timing independent of external sensory cues by incorporating trials with a 3.5-second tone and the same reward window as for the predictable time cue condition (Fig. 3c). This design enabled the distinction between externally driven stimulus-based timing and internally generated temporal estimation mechanisms.

**Figure 3.**
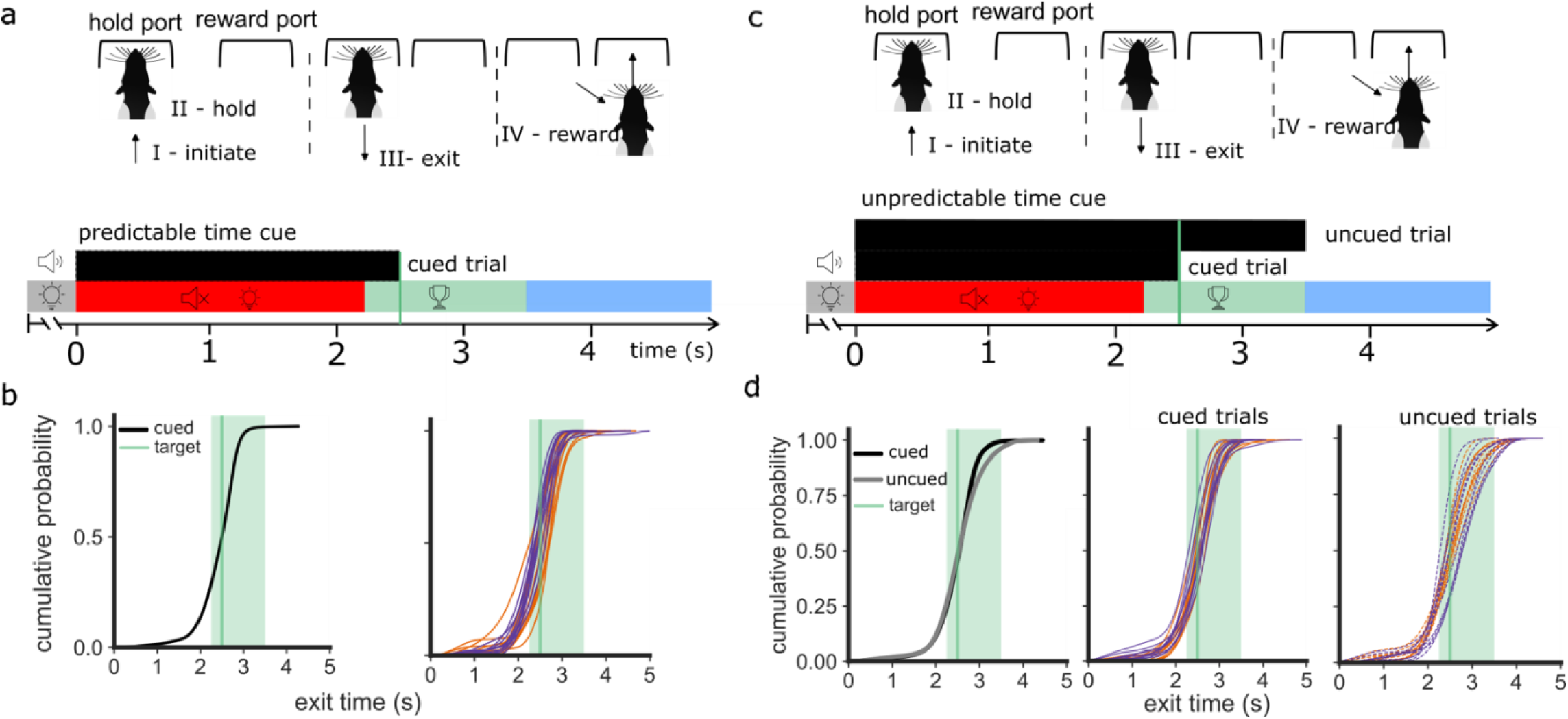
Rats perform an interval timing task. **(a)** Task structure indicating the action sequence (I-IV) **(top**) during a trial in the interval timing task with predictable time cue. The timeline **(bottom)** indicates the trial sequence of possible stimulus events and trial outcomes in the operant chamber. A rat initiates a trial by poking into the hold port (I), triggering, after a random delay, an auditory stimulus of 2.5 s (black bar), which remains the same across the session and therefore the cue becomes predictable, during which the rat fixates its nose into the hold port (II). The trial outcome depends on when the rat exits the hold port, which is measured by the exit time (III). The grey bar indicates the time period in which an exit is too early; the red bar indicates the time period in which an incorrect trial occurs if the rat exits after stimulus onset but before the start of the reward window and results in the sound turned off and stimulus lights on; and the green bar indicates a correct trial in which a reward can be triggered in the reward port (IV). The time it takes for the rat to move from the hold port to the reward port is further referred to as the reward latency. The blue bar indicates the time period of too late trials in which the rat did not exit from the hold port in the reward window. The rat will receive a reward only if it exits the hold port within the reward window (light green). **(b, left)** Cumulative probability of exit times across all animals (n=20 rats) during one interval timing session with predictable time cue. **(b, right)** Individual cumulative probability distributions colour coded by the virus group (EGFP blue and hM4D(Gi) orange, n=10 in each group) during interval timing session with predictable time cue. **(c)** Interval timing task with randomly interleaved cued and uncued trials, in which the action sequence remains the same as in (a) however an unpredictable time cue (upper black bar) which is represented by the longer black bar is also presented in 50 % of the trials. **(d, left)** Exit time cumulative probability distributions across all animals (n=20) during an interval timing session with unpredictable time cue for the cued (2.5s) and uncued (3.5s) trials. **(d, middle)** Individual cumulative probability distributions (n=20 rats) during an interval timing session with unpredictable time cue during cued trials. **(d, right)** Individual response distributions for the uncued trials (n=20 rats).

Rats were first trained on the predictable time cue task (cued trials). The cue consisted of a white noise stimulus. Each trial was initiated by the rat poking its nose into the hold port. After a random delay of 0.5-1.5 s, a sound with a fixed duration of 2.5 s was presented which served as a predictable cue for the rat to exit from the hold port at sound offset (Fig 3a). The rat was rewarded with a food pellet if it exited the hold port during the reward window 2.25-3.5 s from the start of the sound cue. The rats could learn this interval timing behaviour with predictable time cue after approximately 7 days of training. When trained (> 50% accuracy on two consecutive sessions and < 30% too early responses), the distribution of exit times for individual animals in response to the predictable time cue displayed a range of exit times that was centred around 2.5-3 s (Fig. 3b), which closely aligns with the reward window (green shading Fig. 3b), with a mean exit time of 2.42 ± 0.43 s (mean ± SD).

After rats were trained and tested on the interval timing task with the predictable time cue (cued trials), they moved on to sessions where cued trials occurred randomly in 50% of the trials. In the other 50% of trials the same sound was played but for a longer duration (3.5 s Fig. 3c). The latter are referred to as uncued trials because exit from the hold port was not cued by offset of the sound. Uncued trials had the same reward window as that used in cued trials. Therefore, in these sessions, uncued trials tested if rats (based on prior experience and reinforcement), could reliably estimate the time interval from the sound onset to exit the hold port to receive the reward.

After animals were trained on the interval task with both cued and uncued trials (>50% accuracy on two consecutive sessions and <30% too early responses), the mean group exit time for cued trials showed a distribution similar to that observed in the interval timing task with predictable cue trials alone, with a mean exit time of 2.47 ± 0.45 s (mean ± SD) (Fig. 3d), and a similar mean exit time for the uncued trials (2.52 s ± 0.53). For individual animals, the distribution of exit times in response to the cued and uncued trials showed a similar pattern of exit times centred around 2.5-3 s (Fig. 3d). There was no significant difference in the exit times between the two trial types (Mann-Whitney U test, U = 103727.5, p = 0.19). This indicates that the animals could perform the interval timing task for cued and uncued trials with similar success.

### Chemogenetic manipulation during predictable timing delays action initiation

When rats achieved stable performance in the predictable time cue task, CNO or vehicle was injected intraperitoneally on separate sessions with one session per condition (see Methods). First, we assessed if CNO treatment affected task performance (defined as the proportion of correct trials where the rat exited the hold port within the reward window expressed as a percentage of the total number of initiated trials within one session). A generalized linear mixed model (GLMM) was fitted to the data, where reward outcome was the dependent variable and manipulation (vehicle or CNO) and group (EGFP or hM4D(Gi)) were the independent variables. No statistically significant difference was found for the interaction between group and manipulation, indicating that there was no observable effect of CNO on task performance (95% CI [-0.120, 0.254] p= 0.485, Fig. 4a). Next, to assess if CNO treatment affected the time taken for the rat to exit from the hold port a linear mixed model (LMM) model was fitted to the data, with exit time as the dependent variable, manipulation and group as the independent variables and animal ID, trials and session date as the random terms. There was a statistically significant effect of the interaction between group and manipulation, with an average increase of 150 ± 27 ms in exit time for the hM4D(Gi) group following injection of CNO (95% CI [97.45, 204] ms, F= 30.703, p-value < 0.001, Fig 4b). Similarly, there was a statistically significant effect of the interaction between CNO and group on reward latency (Fig. c), with an average increase of 80 ms ± 6 (95% CI [67.57, 90] ms, F = 189.468, p-value < 0.0001) for the hM4D(Gi) group following CNO injection. The total number of trials executed in the operant box was analysed using a GLM with number of trials per session as the dependent variable and group and manipulation as the independent variables to determine if CNO manipulation affects overall ability to perform the task. Results indicated that there was also a statistically significant difference in the total number of trials performed. DREADD rats performed 10 % less trials when given CNO compared to control (95% CI [-0.1961, -0.0093], F-value = 4.64, p = 0.031). In summary, these results indicate that although cerebellar manipulation has no detectable effect on overall success rate in performance of the task, it induces slowing of movement, resulting in delayed exit time and increased reward latency as well as a slight reduction in the number of trials performed.

**Figure 4.**
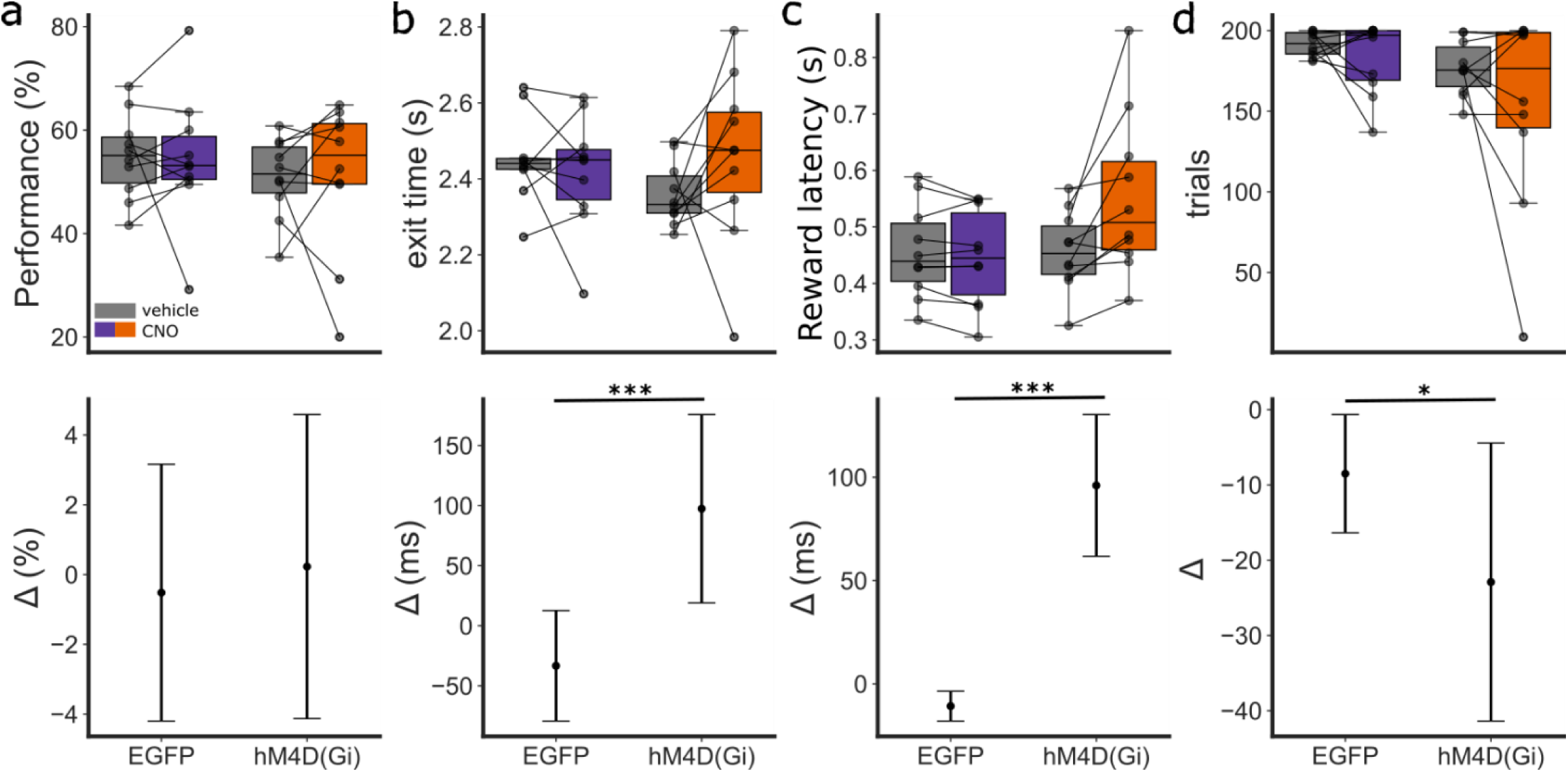
Effect of cerebellar manipulation during interval timing task with predictable time cue. **(a)** Boxplots of the performance showing the group summary statistics (the box shows the middle 50% of the data (interquartile range), with the line inside representing the median. Whiskers extend to the smallest and largest values within 1.5 times the interquartile range) and the distribution of individual mean exit times during one session of the interval timing task with predictable time cue when given CNO (orange, n=10) or vehicle (grey, n=10) in each virus group. Lower plot shows the effect size shown as the average change in performance between vehicle and CNO for EGFP and hM4D(Gi) groups. **(b)** Boxplots and effect size for exit time, plotted in the same way as in (a). **(c)** Boxplot and effect size for the reward latency. **(d)** Boxplots and effect size for the number of trials. *: *p* < 0.01, **: *p* < 0.001 ***: *p* < 0.0001.

### Chemogenetic manipulation during unpredictable timing advances action initiation

The same animals were subsequently exposed to one CNO and one vehicle session (see Methods) in which the tone duration between trials was varied randomly, either with at one duration of 3.5 s(uncued trials), or 2.5 s(cued trials) with the reward time window in both cases occurring between 2.25-3.5 s (as in the predictable time cue task). In each session, the number of cued and uncued trials was balanced and randomized.

To assess if cerebellar manipulation affected task performance, when considering all trial types, a GLMM was fitted to the data, with reward outcome as the dependent variable and manipulation and group as fixed factors. There was a statistically significant effect of the interaction between group and manipulation, with the hM4D(Gi) group performing on average 4.8 % worse than control when given CNO (95% CI-0.379, -0.010], z-value=-2.069, p=0.039, Fig. 5a). To assess if cerebellar manipulation affected the time it takes for the rat to exit from the hold port a LMM was fitted to the data, with exit time as the dependent variable and manipulation and group as the independent variables, with rat ID, trial number and session date as the random term. There was a statistically significant effect of the interaction between group and manipulation, with an average decrease of 107.95 ± 29.33 ms (95% CI [166.60, 49.329], F-value=13.544, p=0.0002) in exit time for the hM4D(Gi) group with CNO (Fig. 5b). There was also a statistically significant effect of the interaction between manipulation and group on reward latency, with an average increase of 6.76 ± 0.61 ms (95% CI [5.562 7.958], F-value = 122.612, p< 0.0001) for the hM4D(Gi) group upon administration of CNO (Fig 5c). The total number of trials executed in the operant box was analysed using GLMM with the total number of trials per session as the dependent variable and group and manipulation as the independent variables and rat ID as the random effect. Results indicated that there was no statistically significant difference in the total number of trials performed (95% CI [-0.102, 0.081], p =0.819, Fig 5d). Overall, these results suggest that cerebellar manipulation during the unpredictable time cue task reduces overall success rate in performance, primarily because rats exit the hold port prematurely.

**Figure 5.**
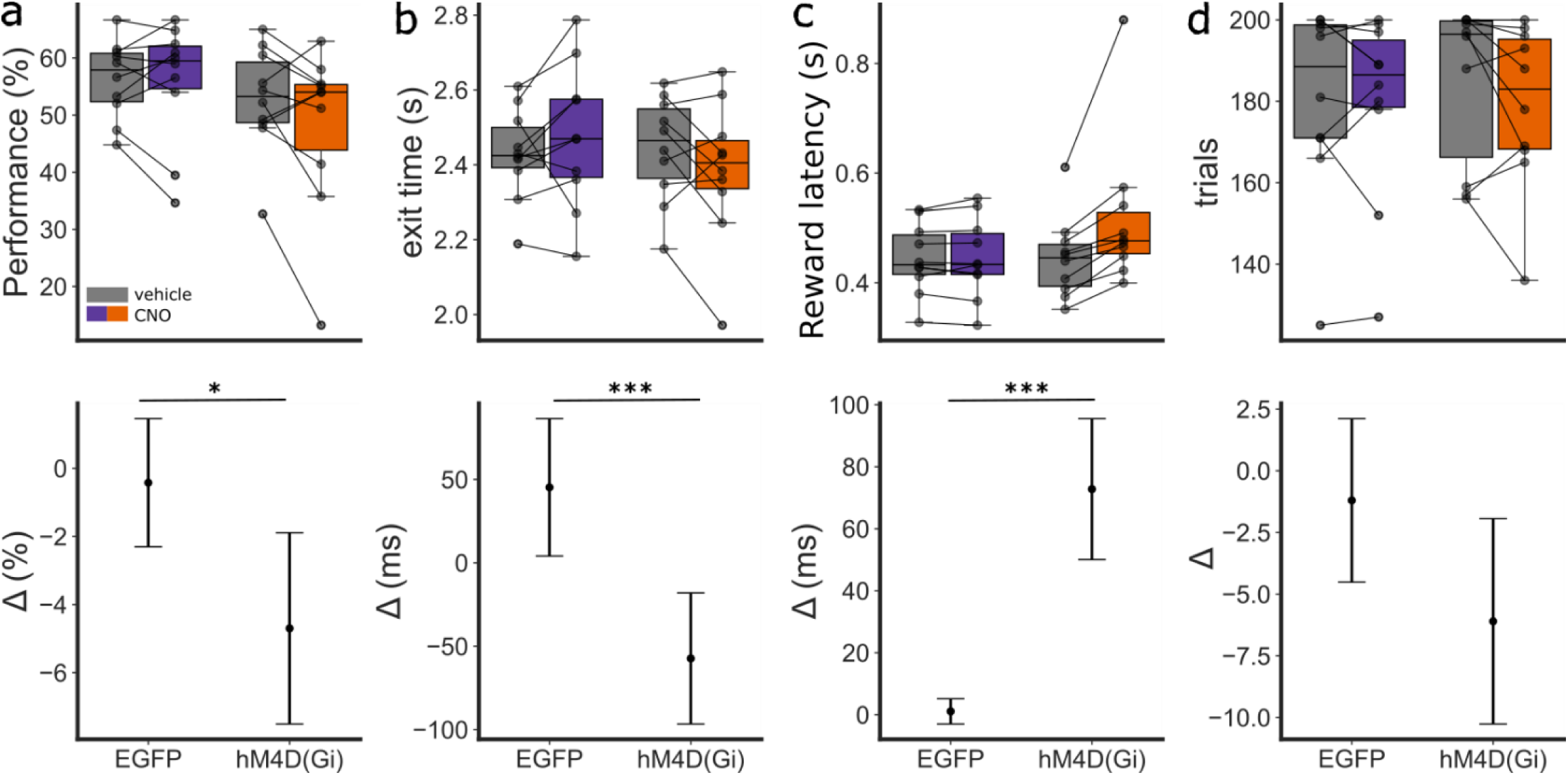
Effect of cerebellar manipulation during interval timing task with unpredictable time cue. **(a)** Boxplots of the performance showing the group summary statistics and the distribution of individual mean exit times during one session of the interval timing task with unpredictable time cue when given CNO (orange, n=10) or vehicle (grey, n=10) in each virus group. **(b)** Boxplots of the exit time plotted in the same way as in a. **(c)** Boxplots and effect size for the reward latency. **(d)** Boxplots and effect size for the number of trials. *: *p* < 0.01, **: *p* < 0.001 ***: *p* < 0.0001.

To distinguish between performance in cued and uncued trials in the unpredictable time cue task, separate analysis of the two trial types was carried out. First, to assess if cerebellar manipulation affects the time it takes for rats to exit from the hold port an LMM was fitted to the data for the cued trial sonly, with exit time as the dependent variable and manipulation and group as the independent variables, with rat ID, trial number and session date as the random term. There was no statistically significant effect of the interaction between manipulation and group on exit time (95% CI [-124.8, 0.002], p= 0.059, Fig. 6Ai and Bi). By contrast, when only uncued trials were considered, there was a statistically significant effect of the interaction between group and manipulation, with an average decrease of 156.44 ± 40 ms (95% CI [-81, -232.15] ms, F-value = 17.0817, p< 0.0001 in exit time for the hM4D(Gi) group with CNO. These results therefore suggest that with the experimental conditions used in the present study, cerebellar manipulation in rats affects internally cued but not externally cued behaviour in a supra-second interval timing task.

**Figure 6.**
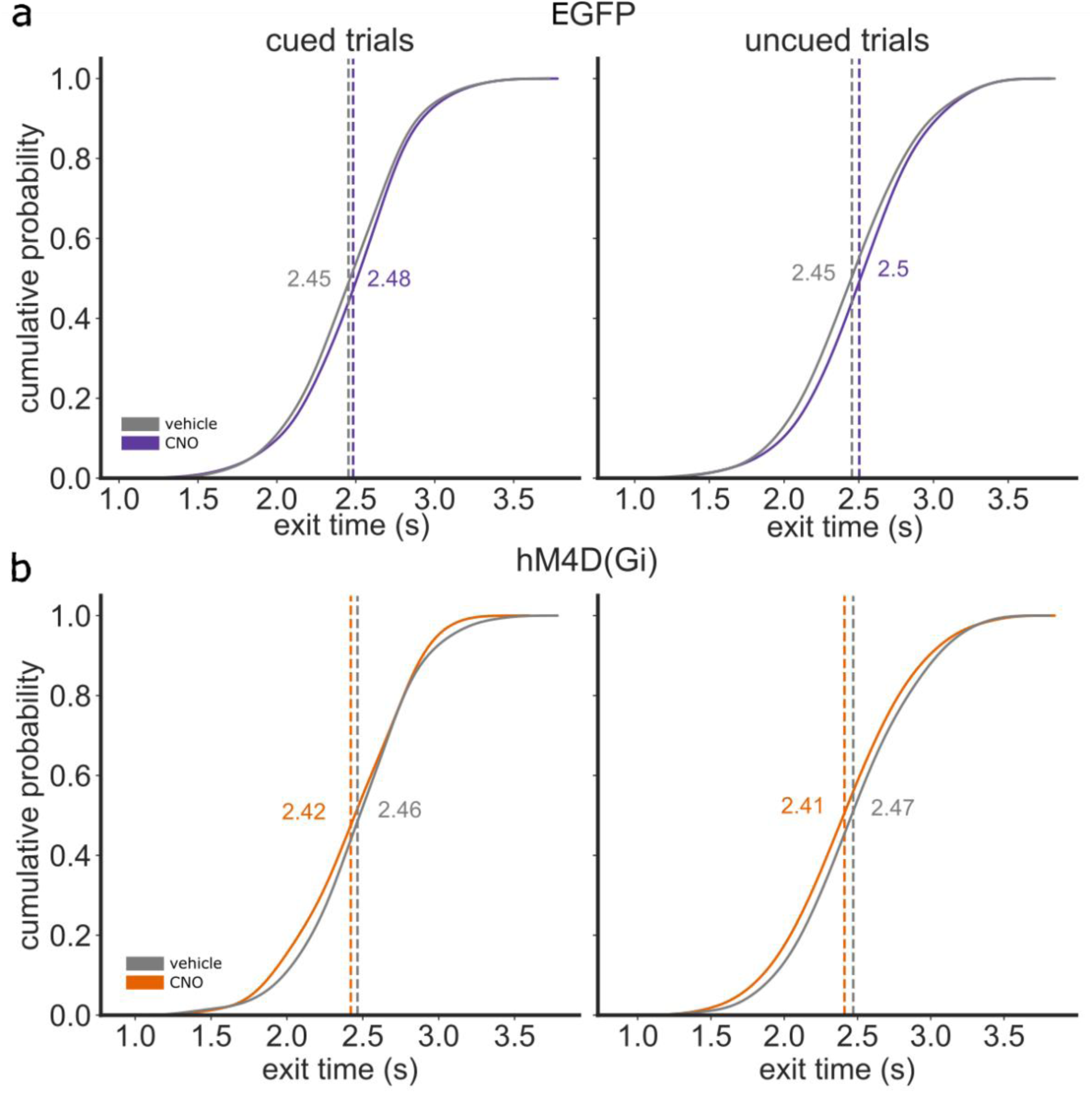
Effect of cerebellar manipulation in cued versus uncued trials. **(a)** Exit time cumulative probability distribution across EGFP animals during a session of the interval timing task with unpredictable time cue under either CNO (orange) or vehicle (grey) conditions. Cued trials (left) occurred randomly in 50% of the trials, and the other 50% of the trials consisted of the uncued trials (right). **(b)** Cumulative probability distribution across hM4D(Gi) animals for cued trials (left) and uncued trials (right). Dotted line indicates the mean exit time as indicated by its respective value.

## Discussion

The current study demonstrates that inhibiting output from the cerebellum, specifically via the LCN, results in behavioural deficits in interval timing in rats. These deficits differ depending on the predictability of the time cue: cerebellar manipulation causes delayed exit from the waiting port when the time cue is predictable, whereas when the time cue was unpredictable, it led to the opposite effect – premature exit times. Given that in the present study the cerebellar manipulation resulted in slowing of locomotion in the open field and slowing of nose poke movement in the predictable and unpredictable cued tasks, it seems reasonable to conclude that the premature exit times were not primarily due to general motor deficits (which would be expected to increase exit time) but rather reflect an impairment in self -timing. While the observed effects on timing behavior were on average a small fraction of one second (0.1-0.15 sec), this may, in part, reflect limitations of the chemogenetic approach, which allows for sustained modulation of neuronal activity rather than precise, real-time inhibition. Nevertheless, even subtle changes in timing precision can be crucial for survival, as many animals rely on accurate temporal estimates to evade predators, optimize foraging, or coordinate social behaviours (Buhusi & Meck, 2005; Coull et al., 2011).

### Comparison with previous studies

Our findings contribute to the ongoing debate regarding the cerebellum’s role in self-timing across sub-second and supra-second intervals. Ohmae et al., (2017) and Kunimatsu et al., (2018) demonstrated that eye movement-related preparatory neural activity in the cerebellar dentate nucleus was temporally consistent across delay intervals, irrespective of duration, suggesting that the cerebellum plays a role in both sub-second and supra-second timing. Specifically, for shorter intervals (<1.2 seconds), the timing of self-initiated saccades was predicted by the rate of increase in dentate neuron firing, with a steeper ramp in activity associated with earlier movement onset. In contrast, for longer intervals (2.4 seconds), preparatory neuronal activity did not begin immediately but instead emerged later in the delay period, and its onset timing correlated with saccade latency. This suggests that for extended delays, the cerebellum adjusts movement timing by modulating when preparatory activity is initiated rather than its rate of increase. Consistent with this, microstimulation of the dentate nucleus during the delay period advanced self-timed saccades by approximately 100 ms, indicating a causal role for the cerebellum in initiating self-timed movements by advancing the onset of preparatory activity (Ohmae et al., (2017; Zhou & Buonomano, 2022; Issa et al., 2020). Conversely, muscimol infusion into the dentate nucleus delayed self-timed saccades at the 2.4-second delay by approximately 100-200 ms and increased trial-by-trial variability in movement initiation, demonstrating that the dentate nucleus is critical for both the timing and precision of self-initiated saccades (Kunimatsu et al., 2018). This aligns with our results, where cerebellar inhibition led to a similar time course to the disruption in self-timing during the unpredictable uncued trials, further supporting a role for the lateral cerebellum in self-timing. The fact that cerebellar inhibition can disrupt timing responses at longer timescales suggests that the cerebellum may not be limited to fine-tuning movement execution but may also contribute to maintaining an internal representation of elapsed time, that includes supra-second timing (Tanaka et al., 2024).

Further supporting our findings, Heskje et al. (2020) demonstrated that pharmacologically inhibiting D1 dopamine receptors in the lateral cerebellar nucleus (LCN) led to increased variability and imprecise timing in a supra-second interval timing task where rats had to estimate elapsed time. Specifically, they observed a decrease in temporal accuracy, with animals responding earlier than required, mirroring the premature responses observed in our unpredictable timing condition following chemogenetic inhibition of the LCN. Furthermore, Heskje et al. (2020) performed electrophysiological recordings in the medial frontal cortex and showed that cerebellar D1 receptor inhibition disrupted ramping activity, a neural signature closely linked to interval timing performance (Buhusi and Meck, 2005; Narayanan, 2016). While our study did not include electrophysiological recordings, the behavioural similarities observed suggest that cerebellar inhibition may impair self-timed responses through disruption of cortical timing mechanisms.

Heslin et al., (2022) investigated in rats the role of the LCN in interval timing but in contrast to our findings did not find any significant effects. However, an important difference with the present study is that Heslin et al., (2022) considered longer time windows, ranging from 4-12 seconds. Another important difference was the type of behaviour studied. While in the present study animals were required to keep their noses within the waiting port and then perform a discrete head movement to obtain a reward, Heslin et al., (2022) used a timing task where the animals were not restricted from carrying out movements such as grooming or pacing while waiting for the fixed interval to lapse. Such behaviours are well known to improve temporal judgements (Fetterman et al., 1998). Nonetheless, the discrepancy with our findings merits further study in which factors such as the time interval employed (seconds versus tens of seconds) and type of movement are taken into account.

Our findings are, however, broadly consistent with results obtained in cerebellar patients (e.g., Gooch et al., 2010). We found that timing impairments were bidirectional based on task demands. Similarly, patients with lateral cerebellar damage show increased errors in time production but decreased errors in time estimation. In the present study one important difference between the predictable and unpredictable time cue tasks is that in the former there is no requirement for the animal to keep track of time, as the stimulus (auditory cue offset) and reward-related response are closely coupled. Whereas in the unpredictable time cue task, stimulus and reward are decoupled (Freestone et al., 2010; Freestone et al., 2013). In terms of reward prediction, the former task can therefore be considered mainly an externally cued response whereas the latter is more an internally cued response.

## Internal models

A key question is how the cerebellum contributes to timing behaviour across different timescales. Traditionally, the cerebellum has been framed within the internal model hypothesis, where it functions as a predictive system that anticipates the sensory consequences of motor actions and fine-tunes movements accordingly (Wolpert et al., 1995; Wolpert et al., 1998; Sokolov et al., 2017) Streng et al., 2022). While internal models have been extensively studied in sub-second motor control, recent evidence suggests that the cerebellum extends its predictive functions to longer timescales, integrating motor and sensory information to form internal representations of elapsed time. For example, cerebellar Purkinje cells have been shown to acquire internal models of supra-second stimulus timing, utilising prediction errors to refine these representations (Narayanan et al., 2024). In zebrafish, cerebellar circuits modulate decision timing and action selection, with cerebellar neurodynamics predicting movement onset more than 10 seconds in advance (Lin et al., 2020). This suggests that cerebellar internal models contribute to long-term motor planning and decision-making. Furthermore, a computational model on cerebro-cerebellar interactions indicates that the cerebellum does not function in isolation but works in concert with the prefrontal cortex, supporting higher-order temporal processing across extended durations (Boven et al., 2023). Specifically, the lateral cerebellar (dentate) nucleus, projects to non-motor cortical areas, suggesting that cerebellar timing computations extend beyond fine motor control to influence cognitive timing mechanisms (Boven & Cerminara, 2023). Additionally, a recent magnetoencephalographic study in healthy participants found that oscillatory cerebellar activity in the beta band (14–30 Hz) is linked to tracking supra-second temporal intervals, reinforcing the cerebellum’s role in proactive motor timing and sensory prediction (Andersen & Dalal, 2024). These findings collectively suggest that the cerebellum’s predictive functions are not strictly limited to movement execution but also involve forming internal representations of time, which are essential for both short- and long-duration behaviours.

In conclusion, the present study provides evidence in rats that the cerebellum’s role in interval timing extends beyond control of movements within sub-second timescales to encompass temporal processing at longer time intervals associated with more complex tasks such as action planning that conventionally are considered to be regulated by higher order structures involved in cognition. By integrating temporal processing across motor and cognitive domains, the cerebellum supports adaptive behaviour in all its forms – from sub-second control involved in conditioned reflexes to supra-second control involved in complex self-timing tasks. Future studies should determine if common cellular processes are involved across this wide temporal range and explore cerebellar interactions with brain-wide neural networks involved in timing and prediction.

## Methods and Materials

### Experimental animals

All animal procedures were performed in accordance with the UK Animals (Scientific Procedures) Act of 1986 under the authority of a UK Home Office Project licence (PPL number PA26B438F) and approved by the University of Bristol Animal Welfare and Ethical Review Body. Experiments were conducted on 20 male Lister Hooded rats (HsdOla:LH, 330-480g at time of surgery, Envigo). Animals were housed in pairs under a 12:12 hour reverse light–dark cycle (light phase 20:15-08:15, target conditions: 20°C and 45–65% humidity). Experiments were therefore performed during the dark phase when they are naturally most active. Food and water were available ad libitum prior to and during recovery from surgical procedures. Water was available ad libitum throughout the experiment. After recovery from surgical procedures (see below) animals were each fed approximately 16 g of standard laboratory chow per day, in addition to food rewards obtained during behavioural tasks. For tasks involving food reward, 45 mg grain-based sweetened reward pellets (TestDiet LabTab AIN-76, 5TUL, catalogue number 1811155) were used. The weights of the animals were monitored 5 days a week to ensure they did not drop below 90% of the normal growth curve. Handling occurred daily one-week prior to surgery and during the recovery phase. Animals were habituated to the experimenter and experimental room for five consecutive days prior to behavioural training. During the experiment animals were handled every weekday.

### Viral vectors

Viral vectors were injected into the lateral cerebellar nucleus (LCN) in order to study the role of cerebellar projections during an interval timing estimation task. Two AAV vectors were used: 10 animals received bilateral injections of DREADD virus (AAV5-hSyn-hM4D(Gi)-mCherry, Addgene, USA, plasmid #50465; titre 1.2×10^12^ gc.ml) whereas 10 animals received bilateral injections of a control virus (AAV5-hSyn-EGFP, Addgene, USA, plasmid#50465; titre 1.2×10^12^ gc.ml). Animals were randomly assigned to the control or the treatment group (hM4Di group). The experimenter was blinded to the identity of the virus that was used for transfection until after the behavioural analysis was complete.

### Surgical procedure

All surgical procedures were performed under aseptic conditions. General anaesthesia was induced by initially administering gaseous isoflurane, followed by an intraperitoneal injection of ketamine (50 mg.kg-1; Vetalar) and medetomidine (0.3mg.kg^-1^; Domitor). Depth of anaesthesia was regularly monitored throughout surgery by testing the hindpaw withdrawal reflex and additional doses of ketamine/medetomidine were given as necessary to maintain surgical anaesthesia. Each animal was placed in a stereotaxic frame with atraumatic ear bars. Throughout the procedure, body temperature was maintained at approximately 37°C with the aid of a thermostatically controlled heated blanket. Eye ointment (LacriLube) was placed on the eyes to prevent corneal injury due to drying. The incision site was treated with local anaesthetic cream (lidocaine). A midline scalp incision was made to access the skull. Bregma and lambda were measured to ensure the skull was level in the dorsoventral plane. Coordinates relative to bregma were measured to allow precise positioning of burr holes for viral injections. The LCN injections were performed bilaterally at AP -11.2mm and ML +/-3.4mm relative to bregma, and DV –4.0mm from the surface of the cerebellum. Virus was delivered using a pulled glass micropipette connected to a 25 µl syringe (Hamilton, Bonaduz, Switzerland) via tubing filled with mineral oil, and was then backfilled with 1 µL of the viral vector using a syringe driver (AL-1000, World Precision Instruments). A volume of 0.5 nl was injected per hemisphere at 200 nl/min and the pipette then left in place for approximately 10 min following viral delivery, to minimise leakage of the virus back up the pipette track. At the end of each surgery, rats were given the medetomidine antidote atipamezole (Antisedan, 0.1 mg intraperitoneally), analgesic (Metacam, 1 mg/kg subcutaneously) and saline (10 ml/kg subcutaneously). Rats were singly housed for 7 days following surgery and then returned to their original pairings. Aminimum period of 6 weeks was allowed for expression of the viral vector before any experimental manipulations. During this period animals underwent behavioural training on the interval time estimation task.

### Interval timing task

The interval time estimation task was performed in a standard light-resistant and sound-proof operant box (Med Associates, OpCoBe Ltd., UK) which were controlled by Klimbic software (Conclusive Solutions LTD, UK). The operant chamber consisted of two nose ports and each port contained a light at the top and a nose poke receptacle with infrared (IR) photobeam at the bottom. The light was used as a reinforcer during the behavioural paradigm. The photobeam served as a head entry detector for nose pokes in each port and was created by a single infrared (IR) light source and receiver. When the beam was uninterrupted, the IR receiver maintained a high output signal, when the rat’s head entered the port the IR beam would be broken, and the receiver would set the out put signal to low. The right port served as the “holdport” in which the rats maintained their head position, while the other port served as the “reward port” for reward collection. Relative positions of the hold and reward port were fixed across the whole experiment. A rubber tube connected the pellet receptacle of the reward port to the pellet dispenser. An audio generator produced tones that were delivered via a speaker placed at the top of the chamber, in between the two ports. The signals from the operant box were recorded through input/ouput cards and interfaced with a computer. Animal behaviour was monitored via a camera (Microsoft LifeCam HD-3000) which was attached to the ceiling of the operant chamber above the hold port. In the interval timing task, food restricted rats learnt to estimate a 2.5 s sound duration through positive reinforcement. Rats indicated the estimated time by exiting from the hold port; if this action was around the target duration of 2.5 s following tone onset, rats could nose poke into the reward port which would trigger delivery of a food pellet. The behavioural protocol used for this experiment is based on the interval timing task described by Xu et al. (2014).

Rats were habituated to the operant box in two successive sessions of 20 minutes per session. Animals underwent training in stages as follows:

1. Tone training: The first stage of the task required the rats to learn to associate a nose poke into the hold port with a 2.5 second auditory cue (white noise, 75 dB), and that a food pellet was delivered in the reward port on termination of the auditory cue. Criterion for the first stage of training was 100 rewarded trials in 30 min on two consecutive days.
2. Action suppression training: Rats then progressed to the second stage where they learnt to actively hold a nose poke for 2.5 s during the auditory cue. The duration of the cue was gradually increased from 0.5 s to the target duration of 2.5 s according to individual performance. Successful holds resulted in a food pellet reward whereas failure to sustain a nose poke lead to a time-out, which consisted of the lights in both the hold and reward ports illuminated for 16 s. Criterion for this second stage was to sustain the nose poke for the required duration for at least 50% of the trials across 2 sessions, with each session consisting of a minimum of 100 trials. All rats learned to initial training stages 1 and two within two weeks.
3. Interval timing training with predictable time cue (Fig. 1A): The next stage of training introduced the reward window and random delay. For each trial, the sound duration was fixed to 2.5 s and served as the cue for the rat to exit from the hold port at sound offset. As the sound offset conveys predictive information for reward availability, this is referred to as a time cue (Freestone et al., 2013). A random delay (drawn from a uniform distribution within 0.5-1.5 s) was introduced between the self-initiated nose poke and sound onset. This required the animal to pay attention to the cue, as the time for reward availability became contingent on the presentation of the cue and not on the nose poke. Exiting the hold port during the random delay was referred to as a “too early” trial and resulted in a time-out. The reward window began 2.25 s after the start of the auditory cue and lasted 1 s after the end of the cue to accurately shape behaviour. Nose poke release during this time triggered a reward pellet and resulted in a “correct trial”. The animals had a duration of 1.5 s to collect their reward following release from the hold port. After reward collection, a new trial could be initiated after an intertrial interval (ITI) of 6 s. Nose poke release that occurred after stimulus onset but before the reward window resulted in an incorrect trial and a time out period. Nose poke releases that occurred beyond the reward window were left unrewarded and considered “too late” trials. Each session lasted 60 mins in which the rats performed 150-200 trials.
4. Interval timing with unpredictable cue (Fig. 1c): After animals were trained and tested on the predictable time cue, animals moved onto the final stage where trials with the predictable time cue of 2.5 s occurred randomly in 50% of the trials, and the other 50% of the trials consisted of the same sound played for a duration of 3.5 s. These trials are referred to as unpredictable trials, because the reward window remained the same and therefore the exit time from the hold port was not cued by the offset of the sound. This stage was designed to assess the rats’ ability to estimate time independently of an external sensory cue, rather than merely anticipate a predictable stimulus. Each session lasted 60 minutes, during which the rats completed 150–200 trials.

### Open field

Open field behavioural testing was performed to determine if chemogenetic inhibition of cerebellar output had any general effect on motor performance. Open field exploration was assessed 30 minutes following CNO injection. Open field testing was separate from behavioural testing on the interval timing task. Rats were placed at the perimeter of a cylindrical arena (90 cm diameter, 51 cm height) which was placed on a black matte plastic base on the floor. Rats were allowed to freely explore the arena for 10 minutes. The behavioural testing occurred in white light (± 140 lux). Behaviour was monitored by an overhead webcam at 30 frames per second (fps). The arena was cleaned with 70% ethanol between each animal.

### CNO administration

CNO administration prior to behavioural testing was performed first on the interval time estimation task with predictable time cue and then on the interval time estimation task with unpredictable time cue. All animals were injected intraperitoneally using a single handed modified restrained method to minimize stress and improve welfare (Stuart et al., 2015). CNO (Tocris, UK) was administered at a dose of 2.5 mg/kg and dissolved in 5% DMSO then diluted in 0.9% NaCl to a final concentration of 2.5 mg/ml. A n e q u i v a l e n t vehicle consisted of 0.9% saline in 5% DMSO. Behavioural testing began 30 minutes following the injection. Typically, testing was carried out over 5 days with day 1, 3 and 5 consisting of baseline sessions and day 2 and 4 consisting of CNO or vehicle administration. The experimenter was blinded to which treatment was administered until after the analysis had taken place.

### Histology and immunostaining

Upon completion of the experiments, animals were anaesthetised with a lethal dose of Euthatal (200 mg/kg Merial Animal Health Ltd, Harlow, UK) and transcardically perfused with 0.9% saline followed by 4% paraformaldehyde. Each brain was dissected and postfixed in 4% paraformaldehyde. After several days, brains were transferred to 30% sucrose in 0.1 M phosphate buffer (PB) and stored until sectioning. Prior to being cut, the cerebellum was removed and embedded in gelatin. A freezing microtome (SM2000R, Leica) using Cryomatrix embedding medium (Thermo Scientific) was used to cut the cerebellum into 40 µm sagittal sections. Sections were collected in 0.01 M PB for preservation and prepared for immunohistochemistry to visualize viral expression of the control or hM4D(Gi) virus. In brief, sections were washed 3 times for 10 minutes in 0.01% PBS to remove debris and placed in 50% ethanol for 30 minutes. After a further 3 x 10 min washes, sections were incubated overnight at room temperature in a chicken anti-eGFP (Abcam ab13970 at 1:2000) or rabbit anti-mCherry (Biovision 5993–100 at 1:2000) anti-body containing 5% normal horse serum. The following day, sections were washed 3 times, for 10 minutes per wash, and incubated for 2h in with secondary antibody (Jackson ImmunoResearch goat anti-chicken Alexa 488 or donkey anti-rabbit Alexa 594 both 1:1000 in PBS-T). Sections were washed 5 min in PBS before mounted on glass slides using 1% gelatin and 0.1% chromium potassium sulphate solution. Fluoromount with DAPI, a stain for all cell nuclei, was applied to the slides before they were cover slipped to prepare for imaging.

### Microscopy

To assess transfection of the virus and placement of the cannula, sections were visualised using a fluorescent Axioskop 2 Plus microscope (Zeiss) fitted with a CoolLED pE-100 excitation system and images acquired using AxioVision software. Transfection of the virus in the LCN was assessed as follows: cerebellar sections were mapped onto standardized sagittal sections of a rat brain using a stereotaxic atlas (Paxinos & Watson, 2009). Sections at key anatomical points 0.18 mm, 0.9 mm, 1.4 mm, 1.9mm, 2.4mm, 2.9mm, 3.4mm, 3.9mm, 4.2mm and 4.6mm from midline were identified and used for manual scoring of fluorescence intensity. Fluorescence intensity was scaled from 0-5, with 0 representing no fluorescence and 5 maximal fluorescence across sections. In each animal, sections with maximum and minimum fluorescence were determined by comparing the sections to each other while keeping illumination settings constant. The section with maximum fluorescence was determined as the section with the largest fluorescent halo around the injection site. Then each section was divided in three regions: cerebellar cortex, nuclei and white matter. A fluorescence score was determined per region by comparing within and across sections. To determine the spread of fluorescence in the mediolateral direction the fluorescence scores of the nuclei were used to visualize the spread of the injection (see figure 2).

### Behavioural measures

#### Open field

A DeepLabCut model (Mathis et al., 2018) was trained to track animals’ movement in the open field using the position of the head. A custom-written Matlab script (version 2021a) was used to extract the total distance travelled as well as the average velocity.

#### Interval timing task

In order to test if chemogenetic manipulation of cerebellar circuits affected the rats’ ability to perform the interval timing task, three metrics were calculated: (i) overall task performance (eq. 1) - the percentage of correct trials, calculated as the number of trials in which the cue was presented and the rat exited the hold port in the reward window, over the total number of trials that the rat initiated, calculated as

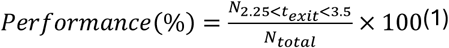

1. (ii) exit time for each trial (eq. 2) - time of hold port release released subtracted by sound onset, both measures with respect to trial start, calculated as

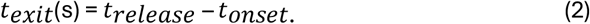

with t(0)=trial start,

and (iii) reward latency for each correct trial (eq. 3) - measured as the time between exit from the hold port and nose poke in the reward port on correct trials, calculated, with t(0)=trial start, as

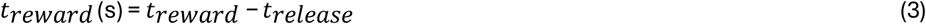

All behavioural measures were extracted from the raw Klimbic file from each session via custom written scripts in Matlab (version 2021a) and Python (version 3.9).

### Statistical Analysis

To assess the effect of CNO on the behavioural metrics of the interval timing task, generalized linear models (GLMs) and linear mixed models (LMMs) were employed. These statistical techniques take into consideration multiple levels of correlation in the dataset (e.g. when collecting multiple trials over different sessions) and were therefore preferred over classic statistical tests such as t-test and ANOVA. LMM were used in the case of normally distributed data whereas GLMMs were used for data following non-normal distributions. LMMs were fit using the lmer function from the lme4 package in R (version 2024.04.2+764). The model included fixed effects for group, manipulation, and their interaction, and random effects for session date, trials and rat to account for repeated measures within sessions and individual differences among rats. The significance of fixed effects was evaluated using Analysis of Variance (ANOVA) with Satterthwaite’s method. GLMs were fit with a binomial distribution and logit link function. For open field analysis, an unpaired t-test was used to compare the control and hM4D(Gi) virus group following a systemic injection of CNO. Data are presented as mean± SD or 95% confidence intervals.

## Acknowledgments

We thank members of the Sensory and Motor Systems group for their feedback, Rachel Bissett, Valentina Pauly and Callum Matthews for their contribution to processing and analysing the histological tissue presented in Figure 2, and Dr. Alex Swainson for assisting with the DeepLabCut models for the open field analysis presented in Figure 3.

## Contributions

EB, JP, RA NLC conceived the project. EB performed the experiments with support from JP. EB performed the data analysis. EB and NLC drafted the manuscript. All authors read and commented on the manuscript.

